# A novel xenograft model reveals invasive mesenchymal transition and ineffective angiogenic response during anti-angiogenic therapy resistance

**DOI:** 10.1101/272328

**Authors:** Arman Jahangiri, William Chen, Ankush Chandra, Alan Nguyen, Garima Yagnik, Jacob Weiss, Kayla J. Wolf, Jung-Ming G. Lin, Soeren Mueller, Jonathan Rick, Maxim Sidorov, Patrick Flanigan, W. Shawn Carbonell, Aaron Diaz, Luke Gilbert, Sanjay Kumar, Manish K. Aghi

**Author notes:** Contributed equally to this work (Arman Jahangiri, William Chen, Ankush Chandra). Corresponding author: Manish K. Aghi, MD, PhD; Professor of Neurological Surgery; UCSF; Diller Cancer Research Building; 1450 Third Street Rm HD-465; San Francisco, CA 94158.

## Abstract

Bevacizumab treatment of glioblastoma is limited by transient responses and acquired resistance. Because of the lengthy duration of treatment that can precede resistance in patients, in order to study changes underlying the evolution of bevacizumab resistance, we created a novel multigenerational xenograft model of acquired bevacizumab resistance. Glioblastoma xenografts were treated with bevacizumab or IgG, and the fastest growing tumor re-implanted into new mice, generating paired isogeneic responsive or resistant multigenerational xenografts. Microarray analysis revealed significant overexpression across generations of the mesenchymal subtype gene signature, paralleling results from patient bevacizumab-resistant glioblastomas (BRGs) that exhibited increasing mesenchymal gene expression correlating with increased bevacizumab treatment duration. Key mesenchymal markers, including YKL-40, CD44, SERPINE1, and TIMP1 were upregulated across generations, with altered morphology, increased invasiveness, and increased neurosphere formation confirmed in later xenograft generations. Interestingly, YKL-40 levels were elevated in serum of patients with bevacizumab-resistant vs. bevacizumab-naïve glioblastomas. Finally, despite upregulation of VEGF-independent pro-angiogenic genes across xenograft generations, immunostaining revealed increased hypoxia and decreased vessel density with increasing generation of treatment, mirroring our findings in patient BRGs and suggesting tumor growth despite effective devascularization caused by VEGF blockade. Besides identifying novel targets for preventing the evolution of resistance and offering a xenograft model for testing resistance inhibitors, our work suggests YKL-40 as a blood biomarker of bevacizumab resistance worthy of further evaluation.

## INTRODUCTION

The premise that the angiogenic switch is critical to malignant tumor progression has been validated by preclinical (1) and clinical (2) studies showing the efficacy of angiogenesis inhibitors. Most angiogenesis inhibitors target vascular endothelial growth factor (VEGF) due to its key role in angiogenesis. Based on encouraging clinical trial results (3), humanized anti-VEGF antibody bevacizumab is approved as monotherapy for recurrent glioblastoma (GBM). Unfortunately, bevacizumab responses are typically transient, with 50% of GBMs (3) that initially respond progressing soon thereafter, with acquired bevacizumab resistance associated with poor outcomes (4). Indeed, phase III trials of bevacizumab in newly diagnosed (5, 6) and recurrent (7) GBM revealed increased progression free survival (PFS) but no impact on overall survival (OS).

While some studies have suggested that resistance to anti-angiogenic therapy is associated with upregulation of compensatory VEGF-independent angiogenesis, others have suggested that resistance involves mesenchymal gene expression changes driving perivascular invasiveness along the remaining blood vessels in an otherwise successfully devascularized tumor (1, 8–10). Furthermore, while xenograft studies have suggested that a short course of tumor treatment with anti-VEGF antibody promotes some degree of mesenchymal transition (11, 12), the impact of duration of treatment, which can be far more prolonged in patients than in xenografts, has yet to be investigated in patients. We hypothesized that the full resistant phenotype emerges gradually over a prolonged course of treatment and involves mesenchymal gene expression changes driving perivascular invasiveness. We investigated this hypothesis in specimens from patients whose tumors progressed after variable duration of bevacizumab treatment and by creating a novel multi-generational xenograft model of bevacizumab resistance.

## RESULTS

To define endpoints for the evolution of bevacizumab resistance that would guide the development of a multi-generational xenograft model, we investigated the impact of bevacizumab-treatment duration on the phenotype of patient bevacizumab-resistant glioblastomas (BRGs). Immunohistochemistry for hypoxia marker CA9 revealed that BRG hypoxia increased with increasing duration of bevacizumab treatment before progression (**Fig. 1a**; P=0.004), while vessel density, assessed by CD31 staining, decreased (**Fig. 1b**; P=0.04), and Ki-67 labeling did not change (**Fig. 1c**; P=0.6). We then used our previously published microarray analysis of 15 bevacizumab-resistant glioblastomas relative to paired pre-treatment glioblastomas from the same patients (10, 13) to analyze expression of the genes previously used to define three glioblastoma subtypes (proneural, mesenchymal, and proliferative) (14). This analysis revealed a significant tendency for tumors to become more mesenchymal and less proneural with increasing duration of bevacizumab treatment before progression (**Fig. 1d**; P=0.01). To determine whether these changes were arising homogeneously in individual cells or instead arising in clusters of single cells, we performed single-cell RNA sequencing of 857 cells from a BRG (**Fig. 2a**). When focusing on the tumor cells (**Fig. 2b**), and moving into the gene space of the 500 most differentially expressed identified by our previously published microarray analysis of 16 BRGs compared to before treatment (10), we identified clusters of single cells (**Figs. 2c-d**), revealing that BRG cells differ in the expression of these genes.

**Figure 1.**
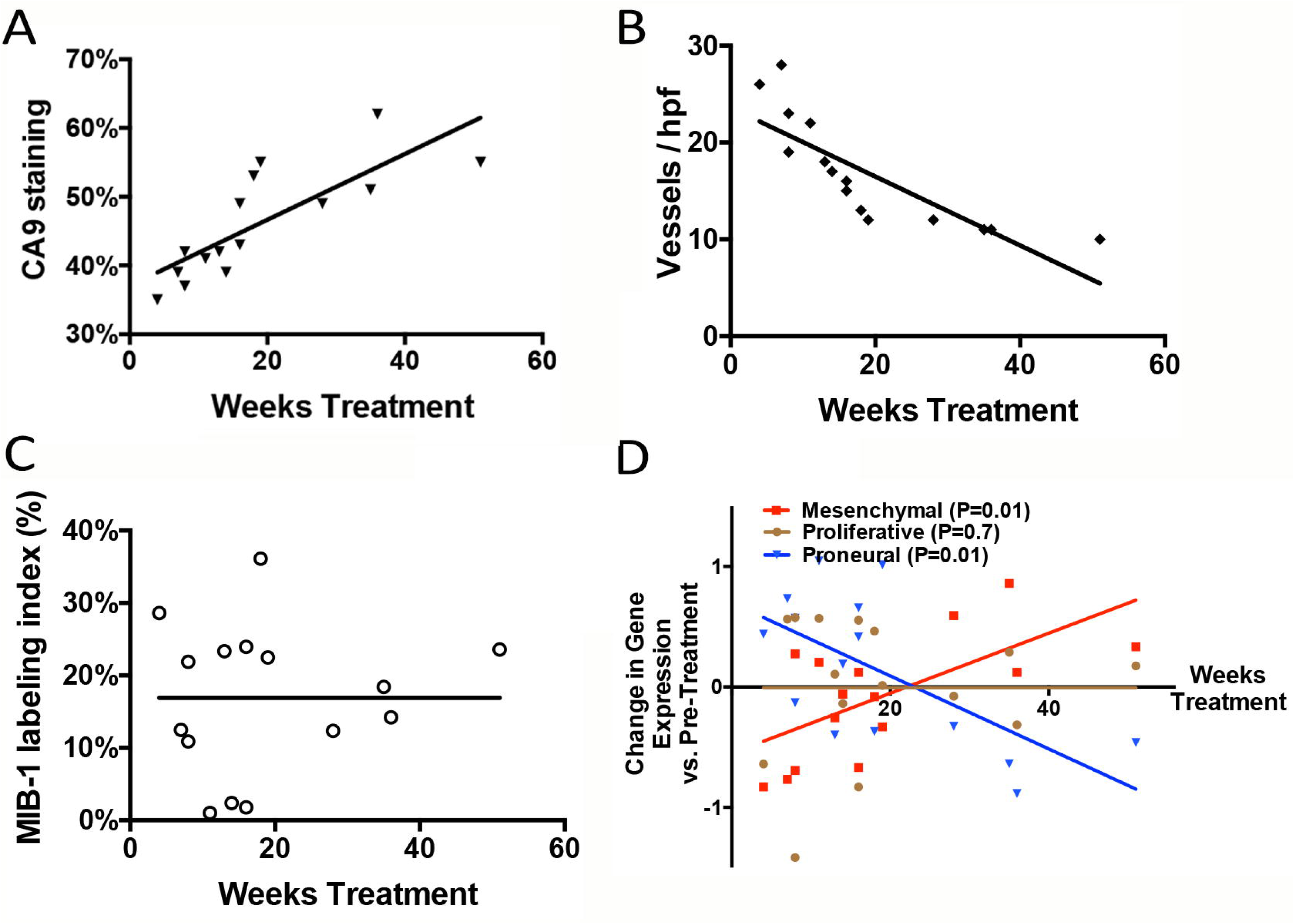
Increasing treatment duration drives hypoxia and mesenchymal gene expression in patient bevacizumab-resistant glioblastoma. (**a**) increased CA9 staining, (**b**) decreased vessel density, (**c**) unchanged proliferation, and (**d**) increased mesenchymal / decreased proneural gene expression were seen with increased duration of bevacizumab treatment in patient bevacizumab-resistant glioblastomas.

**Figure 2.**
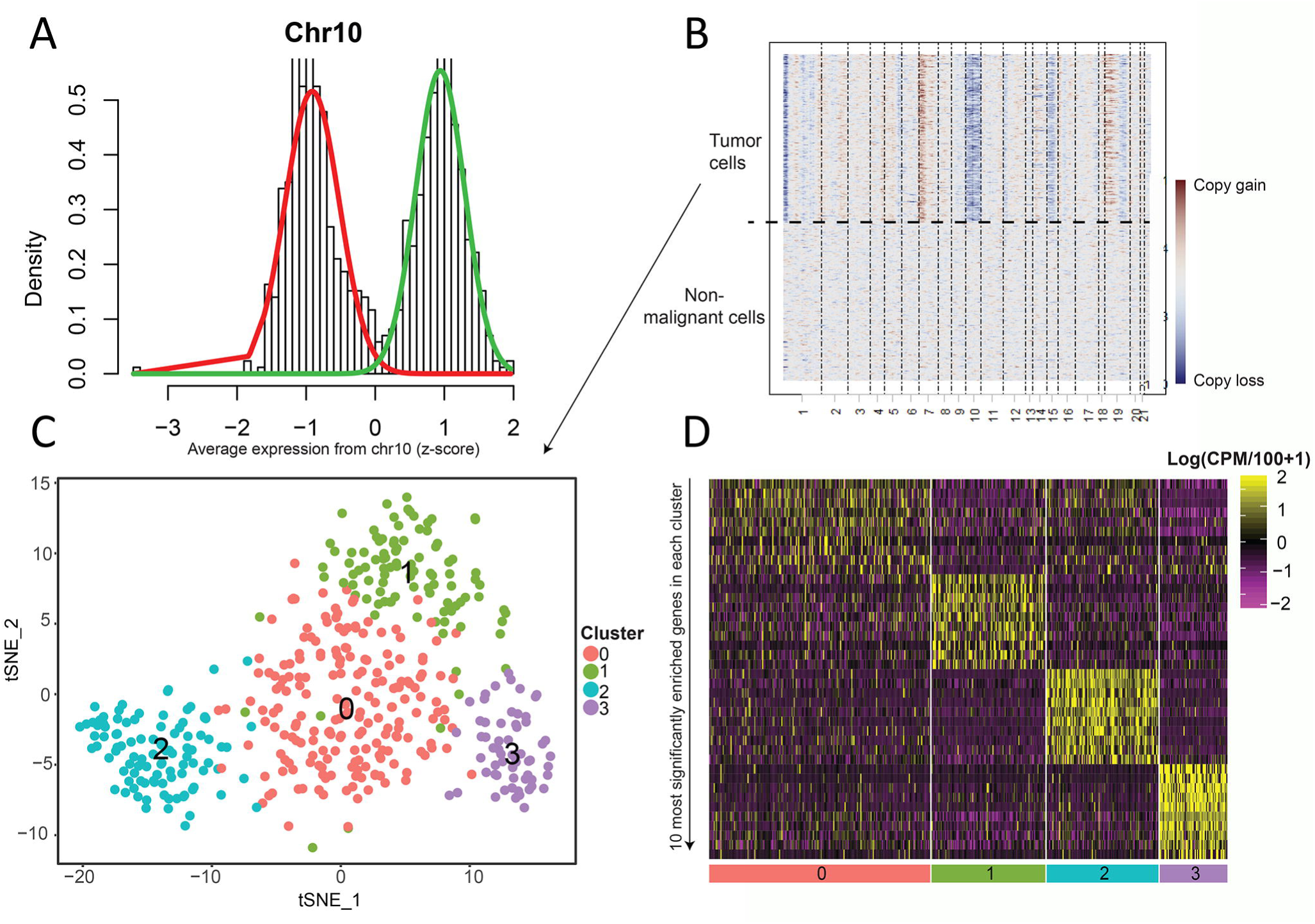
Identification of clusters of single cells from patient bevacizumab-resistant glioblastoma. (**a**) Histogram visualizing the density of cells (y-axis) based on the average expression from chromosome 10 (x-axis). The two components of a gaussian mixture model are indicated by color. The model clearly allows to separate cells into two populations. (**b**) Heatmap visualizing the average expression of genes along chromosomes (x-axis) for all cells (y-axis). Non-malignant cells (bottom) show no copy number alterations, while malignant cells harbor multiple large-scale CNVs, including glioma typical gain of chr7 and loss of chr10. (**c**) Tnse map of single tumor cells (dots). To generate the dimensionality reduced representation, only the 500 genes most strongly differentially expressed between bevacizumab resistant and pre-resistance samples of GBM cases from our previous study were used. (**d**) Heatmap of the 10 most significantly enriched genes (rows) in all cells in each cluster (columns).

To further define changes associated with the evolution of bevacizumab resistance, we established a novel xenograft model of anti-angiogenic therapy resistance (10, 15). Because acquired anti-angiogenic therapy resistance arises after prolonged treatment, U87 xenografts were treated with bevacizumab or IgG during serial *in vivo* subcutaneous passaging for 9 generations, with the most resistant xenograft selected for repropagation each generation (**Fig. 3a)**, creating 9 generations of U87-Bev^R^ and U87-Bev^S^ xenografts, whose bevacizumab-resistance versus responsiveness, respectively, were confirmed in subcutaneous and intracranial microenvironments (10, 15). The ability of the model to replicate human bevacizumab-resistant glioblastomas and the necessity of multiple generations of treatment to model patient resistance was confirmed because four key features in bevacizumab-resistant glioblastomas (4, 13) appeared in the xenograft model but did not culminate until the fifth generation of U87-Bev^R^. First, resistance rate, the frequency of tumors reaching volumetric endpoint by 45 days, rose from 13% to 100% during the first 5 generations (**Fig. 3b; Supplementary Figs. S1-3**). Second, tumor growth rate increased with each U87-Bev^R^ generation until surpassing that of U87-Bev^S^ by the fifth generation (**Fig. 3c; Supplementary Figs. S1-3**). Third, the tumors exhibited decreased vessel density (**Fig. 3d**) and increased hypoxia (**Fig. 3e**) with increasing generation of treatment. Fourth, Ki67-labeling did not change with each generation of treatment (**Fig. 3f**). Thus, like patient BRGs, our multi-generational xenograft model exhibited increased hypoxia with decreased vessel density with increasing duration of bevacizumab treatment, suggesting that bevacizumab-resistant glioblastomas grow despite successful bevacizumab-induced devascularization.

**Figure 3.**
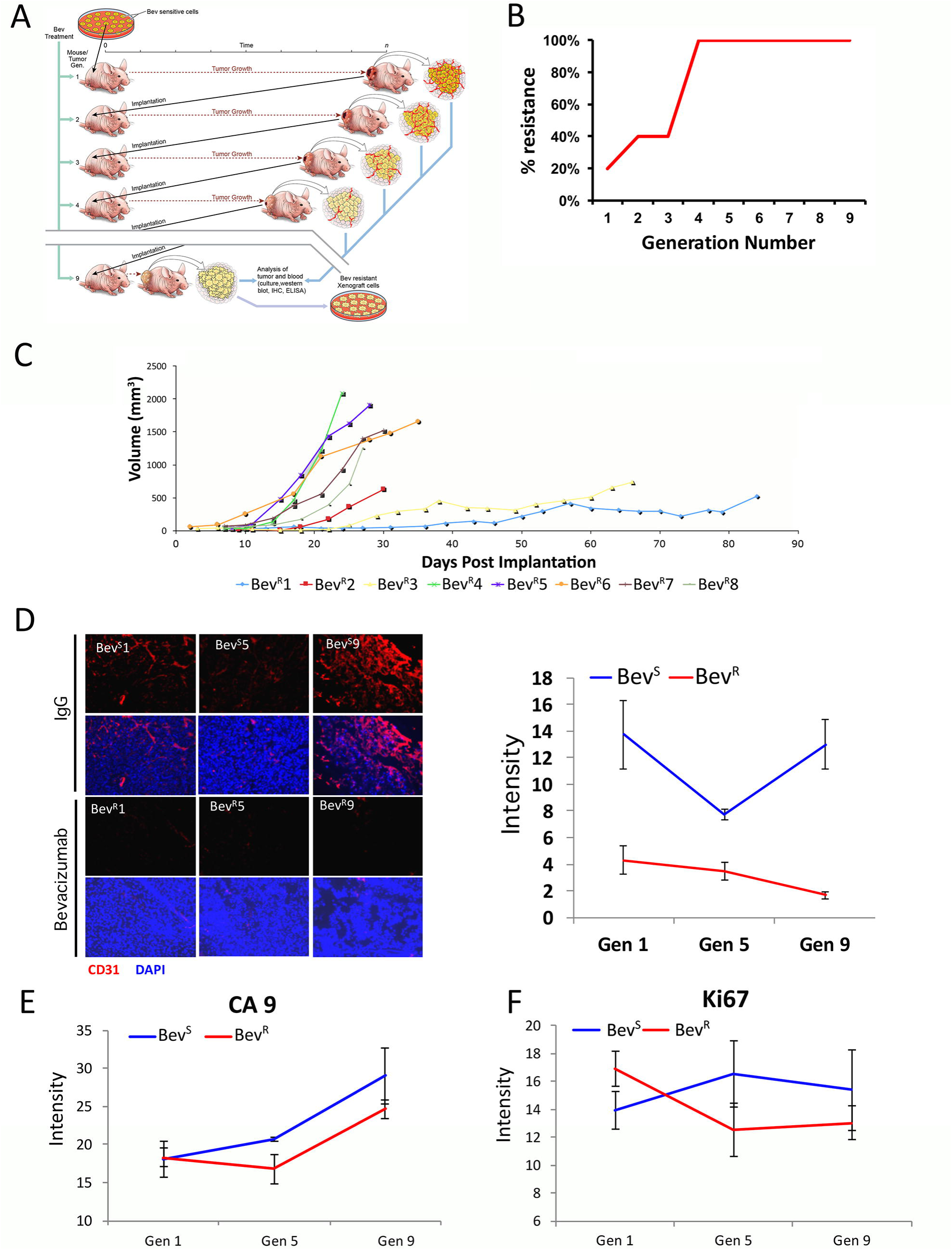
A multi-generational xenograft model replicates the phenotype and gene expression of patient bevacizumab-resistant glioblastoma. (**a**) Schema of the approach used to create the multi-generational bevacizumab-resistant xenograft model. (**b**) The percentage of tumors that were resistant to bevacizumab increased with each U87-Bev^R^ generation. (**c**) U87-Bev^R^ tumor growth rate increased with each generation. (**d**) Decreased vessel density was noted with increasing generation of U87-Bev^R^. (**e**) increased hypoxia density was noted with increasing generation of U87-Bev^R^. (**f**) MIB-1 labeling did not change with each generation of U87-Bev^R^.

To serially investigate transcriptional changes associated with the evolution of bevacizumab resistance, we performed microarray expression analysis of U87-Bev^R^ generations 1, 4, and 9, with generation four of U87-Bev^R^ chosen because of the dramatic increase in tumor growth rate that occurred during this generation (**Fig. 3c**). Gene set enrichment and ontology analysis revealed significant upregulation across U87-Bev^R^ generations 4 (**Fig. 4a**) and 9 (**Fig. 4b**) of genes involved in metabolism, angiogenesis, and inflammatory response, and downregulation of genes involved in DNA packaging and translation, chromatin assembly, and mitosis.

**Figure 4.**
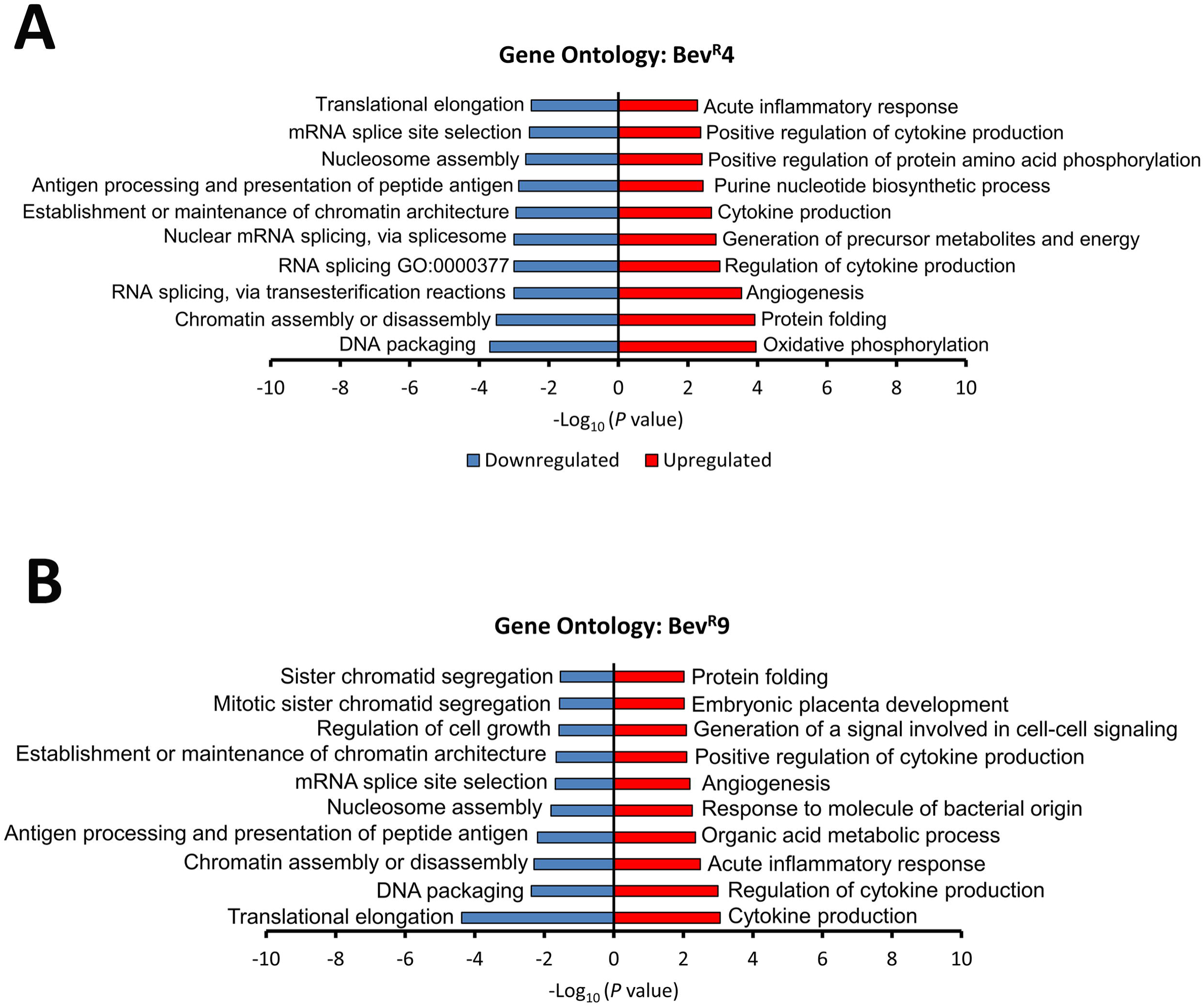
Significantly altered gene ontologies during evolution of bevacizumab resistance in a multi-generational xenograft model. GAGE gene set enrichment analysis revealed enrichment and upregulation of gene ontologies associated with metabolism, angiogenesis, and inflammatory response, and downregulation of genes involved in DNA packaging and translation, chromatin assembly, and mitosis. The top ten enriched gene ontologies are displayed.

To determine if our multigenerational xenograft model replicated the mesenchymal gene expression changes we noted in bevacizumab-resistant patient glioblastomas, we investigated the set of U87-Bev^R^ genes significantly dysregulated across xenograft generations, finding that mesenchymal genes were significantly over-represented and upregulated, while proneural genes were significantly under-represented and downregulated (**Fig. 5a-b**). To validate these findings, we performed qPCR for 14 genes in our array that were also part of a previously described mesenchymal gene set in GBM (14) and found increased expression in all but one gene by generation 9 (**Fig. 5c**). To integrate and elaborate upon these findings, we averaged the expression of these 14 mesenchymal genes by qPCR, along with the 15 proneural and 5 proliferative genes described in the same study (14) that were also part of our microarray to obtain mesenchymal, proneural, and proliferative gene expression scores on a +1 to −1 scale. Doing so revealed increased mesenchymal gene expression approaching +1 by generation 4 and persisting thereafter (**Fig. 5d**), mirroring mesenchymal gene expression changes which occurred with increasing duration of bevacizumab treatment in patient BRGs. Finally, to determine if the secretion of any of these factors in the setting of bevacizumab resistance was high enough to elevate them in the serum and serve as a potential biomarker of this bevacizumab-induced mesenchymal change, we investigated levels of YKL-40 in the serum of patients with BRGs (n=8) versus bevacizumab-naïve GBMs (n=11), finding significantly elevated serum levels of YKL-40 in BRG patient serum compared to serum from recurrent bevacizumab-naïve GBMs (P<0.05; **Fig. 5e**).

**Figure 5.**
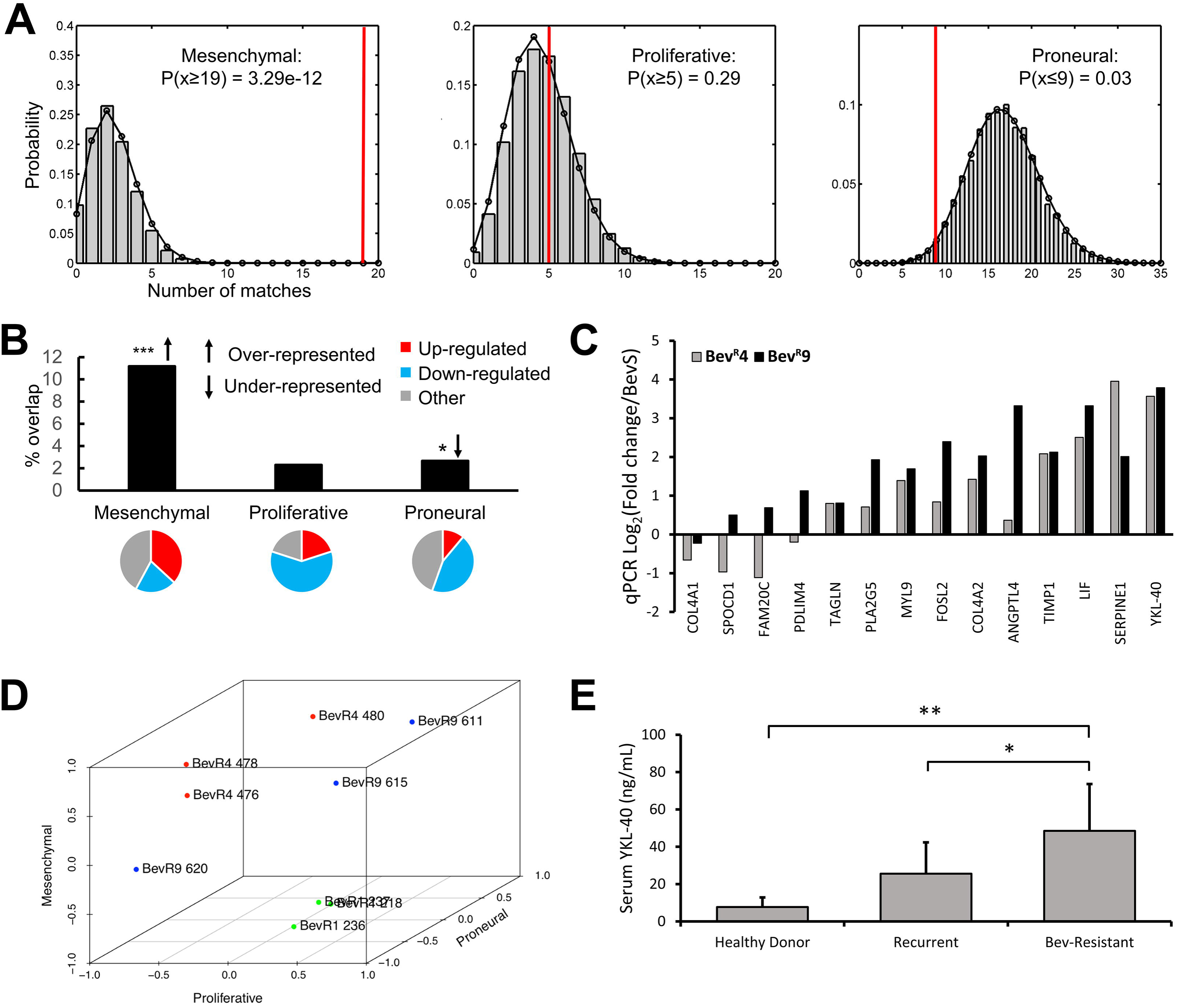
A multi-generational xenograft model replicates the mesenchymal gene expression of bevacizumab-resistant glioblastoma. (**a-b**) Bootstrapping analysis revealed the Philips et al mesenchymal gene set to be highly over-represented and proneural gene set to be under-represented among genes significantly dysregulated across U87-Bev^R^ generations, with 19 of 170 (11%) mesenchymal genes and 9 of 338 (2%) proneural genes found to be dysregulated. 37% of dysregulated mesenchymal genes were upregulated, while 44% of dysregulated proneural genes were downregulated. (**c**) Increased expression of signature mesenchymal genes was confirmed by qPCR, with marked upregulation of the secreted factor, YKL-40. (**d**) Expression of signature mesenchymal, proneural, and proliferative genes were normalized and plotted on a −1 to +1 scale, revealing the evolutionary trajectory of transcriptional subtypes across U87-Bev^R^ generations, with increased mesenchymal gene expression approaching +1 by generation 4 and persisting thereafter. (**e**) ELISA reveals elevated YKL-40 levels in patients with GBMs resistant to bevacizumab compared to patients with recurrent bevacizumab-naïve GBM and healthy donors *, p<0.05, **, p<0.01, ***, p<0.001

Because our microarray analysis revealed multiple dysregulated genes implicated in cytoskeletal and extracellular matrix re-organization and regulation of tumor cell migration and invasion, including increased expression of RHOQ and RHOE, FYN, FGD3, TIMP1, SERPINE1, HBEGF, DUSP1 as well as decreased expression of a tumor motility suppressor, DPYSL3 (**Fig. 6a**), we investigated whether these mesenchymal and invasive gene expression changes noted in U87-Bev^R^ were associated with the altered morphology and increased invasiveness that we have previously described in BRGs (4, 10, 13, 15) and which are hallmarks of mesenchymal cancers in general. Consistently, U87-Bev^R^ cells exhibited more stellate morphology (P<0.01; **Fig. 6b-c**) and increased invasiveness in Matrigel assays compared to U87-Bev^S^ cells (P=0.003-0.04; **Fig. 6d; Supplementary Fig. S4**). We then performed further investigation of the invasive potential of U87-Bev^R^ and U87-Bev^S^ cells in three-dimensional bioengineered models. First, we studied these cells in hydrogel (16) platforms that modeled the gray and white matter along which peritumoral invasion occurs and contained the hyaluronic acid shown to be upregulated in the extracellular matrix of bevacizumab-resistant tumors (17). In these hydrogel platforms, U87-BevR cells proved more invasive than U87-BevS cells (P<0.001; **Fig. 6e**). Then, we studied these cells in microchannel platforms containing artificial blood vessels to model perivascular invasion and found U87-Bev^R^ cells to also be more invasive than U87-Bev^S^ cells in this context (P<0.05; **Fig. 6f; Supplementary Figs. S5-6**).

**Figure 6.**
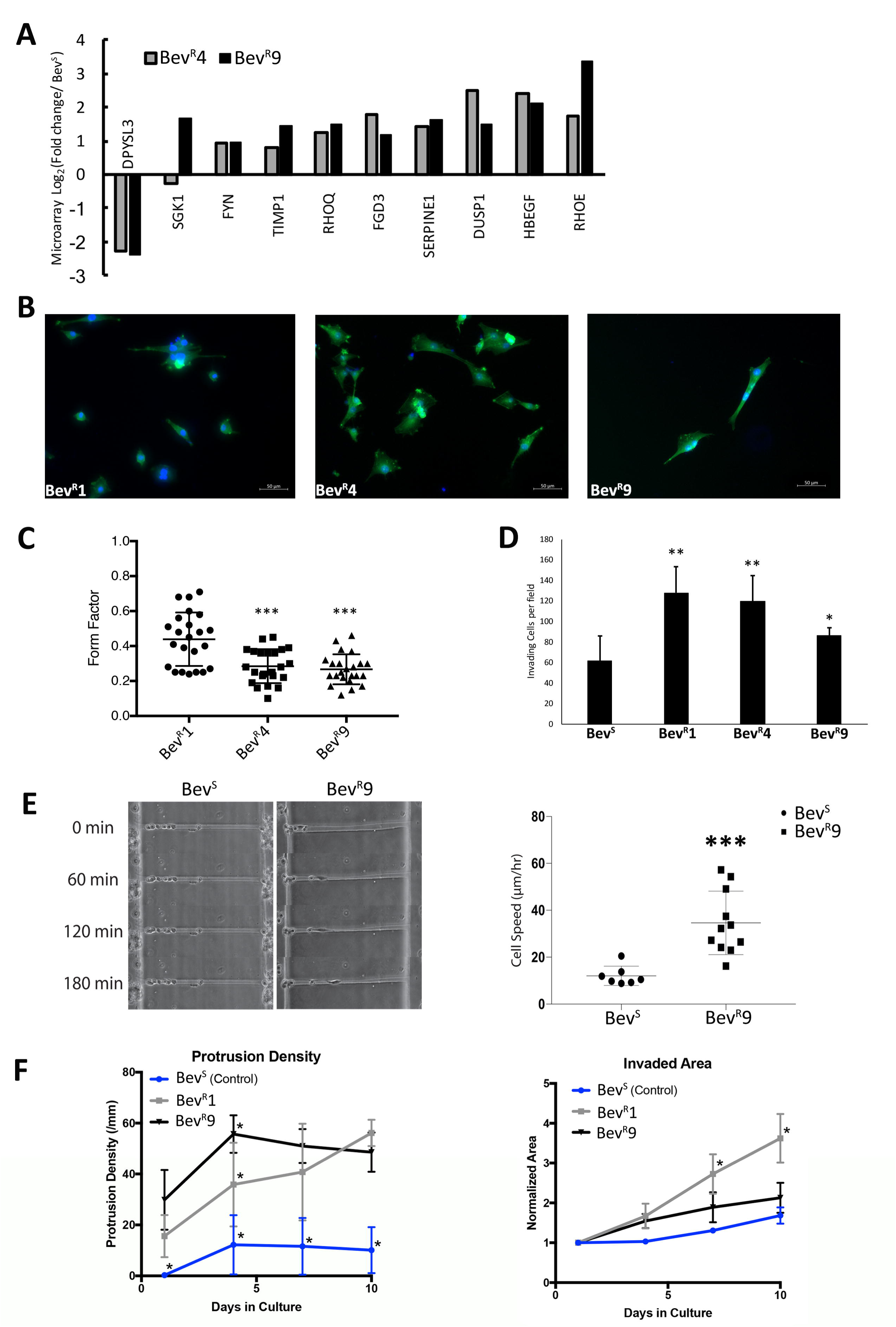
A multi-generational xenograft model replicates the invasiveness of patient bevacizumab-resistant glioblastomas. (**a**) Microarray analysis revealed dysregulation of multiple genes in U87-Bev^R^ (versus U87-Bev^S^) associated with cytoskeletal and extracellular matrix re-organization, migration, and invasion. (**b**) Immunofluorescent imaging of cells from generations 1,4 and 9 of U87-Bev^R^ with Phalloidin reveals cellular morphology. (**c**) Morphological analysis of generations 1,4 and 9 of U87-Bev^R^ reveals lower form factor associated with generations 4 and 9 of U87-Bev^R^. (**d**) Matrigel invasion assay reveals a higher invasiveness of multiple generations of U87-Bev^R^ when compared to U87-Bev^S^. (**e**) 3D bioengineered hydrogel assay modeling white matter tracts and (**f**) 3D bioengineered microchannel platform modeling perivascular invasion through hyaluronic acid (HA) matrix reveal a higher invasiveness (P<0.001 hydrogel and P <0.05 microchannel) of multiple generations of U87-Bev^R^ compared to U87-Bev^S^ cells. *, p<0.05, **, p<0.01, ***, p<0.001. Scale bar= 50μm

Our transcriptional analysis also identified significantly dysregulated genes previously shown to be markers of or to be involved in regulation of glioma stem cells, including CD44, LIF, IL6, and ZEB1, leading us to investigate whether patient BRGs exhibited enrichment of a tumor stem cell population. Prospective plating of cells from patient BRGs (n=4) vs. bevacizumab-naïve recurrent glioblastomas (n=3) in neurosphere medium revealed significantly increased cell counts from dissociated neurospheres in BRGs compared to bevacizumab-naïve GBMs (**Fig. 7a**; P<0.001). Similar culture of U87-Bev^R^ cells in neurosphere medium revealed that, although U87-Bev^R^ cells formed fewer neurospheres than U87-Bev^S^ cells, U87-Bev^R^ neurospheres were larger (p=0.002) and markedly more cellular (p=0.0002) as compared to U87-Bev^S^ cells (**Fig. 7b-f**), reflecting less differentiated, more proliferative stem cells with greater resistance to hypoxia (18). To investigate potential mediators of the increased stem-cell like behavior and mesenchymal change identified in U87-Bev^R^ xenografts and BRGs, qPCR was performed on explanted U87-Bev^R^ and U87-Bev^S^ tumors for GLUT3 and ZEB1, genes associated with both properties (19, 20), revealing increased GLUT3 and ZEB1 gene expression in U87-Bev^R^ xenografts (P=1X10^−5^-0.02; **Fig. 7g**). We then investigated the role of ZEB1 in driving the mesenchymal gene expression changes we had identified in BRGs and our xenograft model. Using CRISPR to disrupt ZEB1 expression in generations 1, 4, and 9 of U87-Bev^R^ and U87-Bev^S^ (**Supplementary Fig. S7**) caused complete loss of the mesechymal gene expression we had seen in U87-Bev^R^ (**Fig. 7h**).

**Figure 7.**
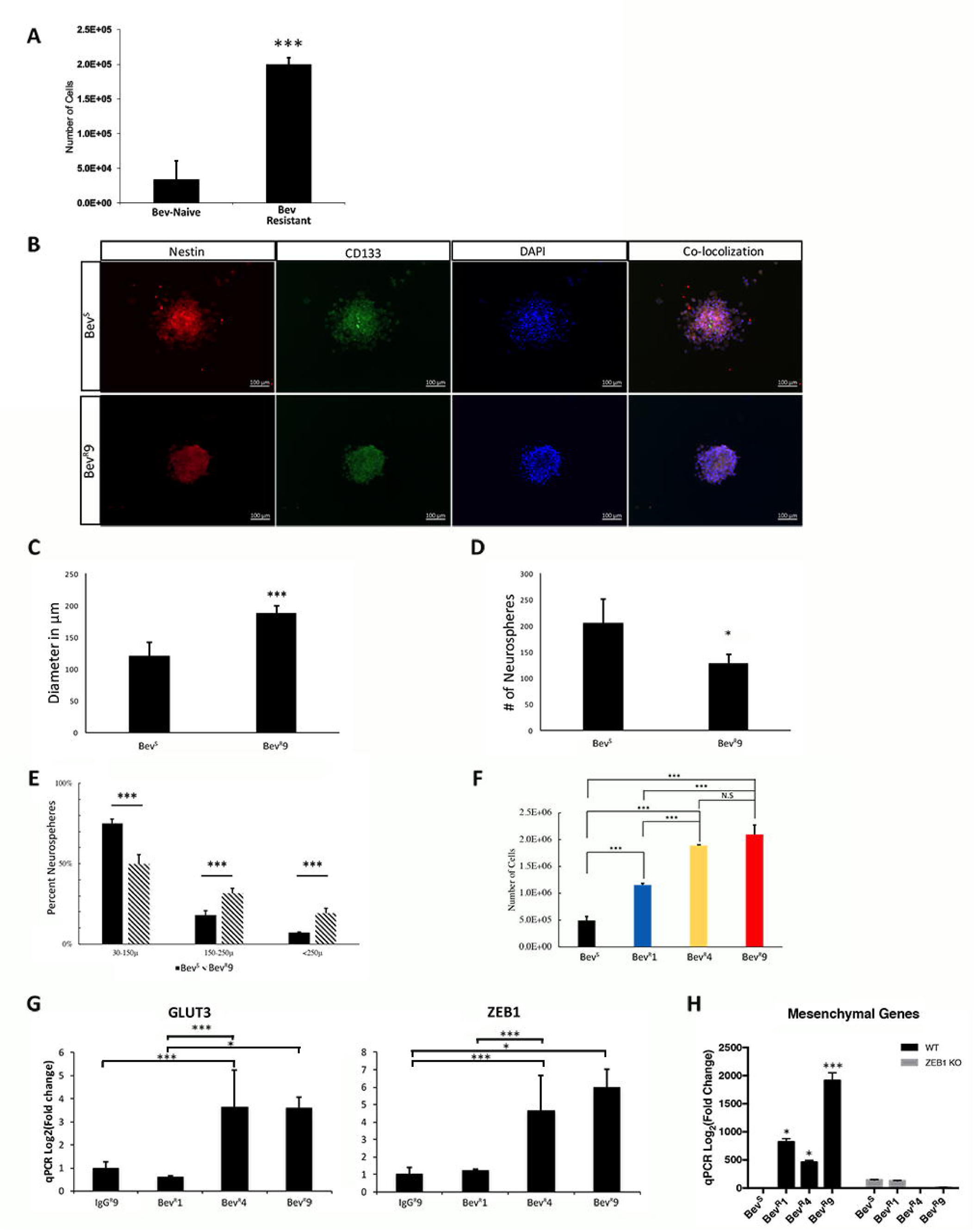
A multi-generational xenograft model replicates the stem cell-enrichment of bevacizumab-resistant glioblastomas. (**a**) Higher total stem cell count yield from neurospheres derived from bevacizumab-resistant GBMs (n=4) when compared to bevacizumab-naïve recurrent GBMs (n=3) (p<0.001). (**b**) Immunofluorescent staining of neurospheres derived from U87-Bev^s^ and U87-Bev^R^ generation 9 cells was done with Nestin and CD133 (stem cell markers). (**c-d**) Neurosphere formation assay revealed a much larger diameter of U87-Bev^R^ generation 9 neurospheres (p=0.002), while Bev^S^ cells yielded a larger number of neurospheres (p=0.03). (**e**) Relative distribution of Bev^S^ and Bev^R^ generation 9 neurospheres by diameter size. (**f**) Absolute cell counts from U87-Bev^S^ and U87-Bev^R^ generation 9 neurospheres. (**g**) Transcriptional analysis of prominent glioma stemness markers, GLUT3 and ZEB1, of tumor chunks from U87-Bev^S^ generation 9, U87-Bev^R^1,4 and 9 reveals a positive correlation between generation number and GLUT3 and ZEB1 mRNA expression in U87-Bev^R^. For ZEB1, U87-Bev^S^ vs U87-Bev^R^ generation 4 and U87-Bev^S^ vs U87-Bev^R^ generation 9, p<0.001 and 0.02, respectively. For GLUT3, U87-Bev^S^ vs U87-Bev^R^ generation 4, p=0.006. (**h**) ZEB1 knockout in U87-Bev^S^ and U87-Bev^R^ generations 1,4 and 9 using CRISPR reveals a significantly reduced transcriptional expression of mesenchymal genes as compared to the wild-type controls of the cell lines. *, p<0.05, **, p<0.01, ***, p<0.001. Scale bar= 100μm.

Interestingly, our microarray analysis also revealed expression changes in angiogenesis pathways, suggesting that anti-angiogenic therapy triggered attempted compensatory upregulation of non-VEGF angiogenic pathways (**Fig. 8a**). Thirty of the most upregulated genes from the significantly enriched angiogenesis gene ontology were selected for qPCR validation, revealing significant overexpression of angiogenesis related genes by generation 1 of U87-Bev^R^ which peaked by generation 4. (**Figs. 8b-c**). Among these upregulated genes were multiple secreted pro-angiogenic factors, including GREM1, FGF2, YKL-40, WNT5A, IL6, IL1A, and IL1B (**Fig. 8d**). However, our immunostaining of xenografts and patient specimens would suggest that this upregulation of genes mediating VEGF-independent angiogenesis failed to meaningfully impact tumor vascularity, based on the reduced vessel density and increased hypoxia noted with increasing treatment duration in resistant patient glioblastomas and xenografts. To determine if this ineffective compensation reflected the failure of upregulated secreted angiogenic genes in U87-Bev^R^ to achieve biologically meaningful levels, we analyzed the effect of conditioned media (CM) from cultured U87-Bev^R^ and U87-Bev^S^ cells on the formation of endothelial tubules in HUVEC cells. We found that U87-Bev^R^ CM drove increased endothelial segment and mesh formation compared to U87-Bev^S^ conditioned media (P=0.0007-0.01) and that, while bevacizumab inhibited U87-Bev^S^ effects on HUVEC cells (P=0.0004), by generation nine, bevacizumab failed to block the increased U87-Bev^R^ effects on HUVEC cells (**Fig. 8e; Supplementary Fig. S8**). Thus, while compensatory pro-angiogenic factors secreted by U87-Bev^R^ cells were able to stimulate a VEGF-independent angiogenic response in culture, the sustained hypovascularity and hypoxia we identified in U87-Bev^R^ xenograft tumors and BRGs may reflect additional *in vivo* effects rendering this compensatory angiogenic response ineffective in the setting of prolonged bevacizumab treatment.

**Figure 8.**
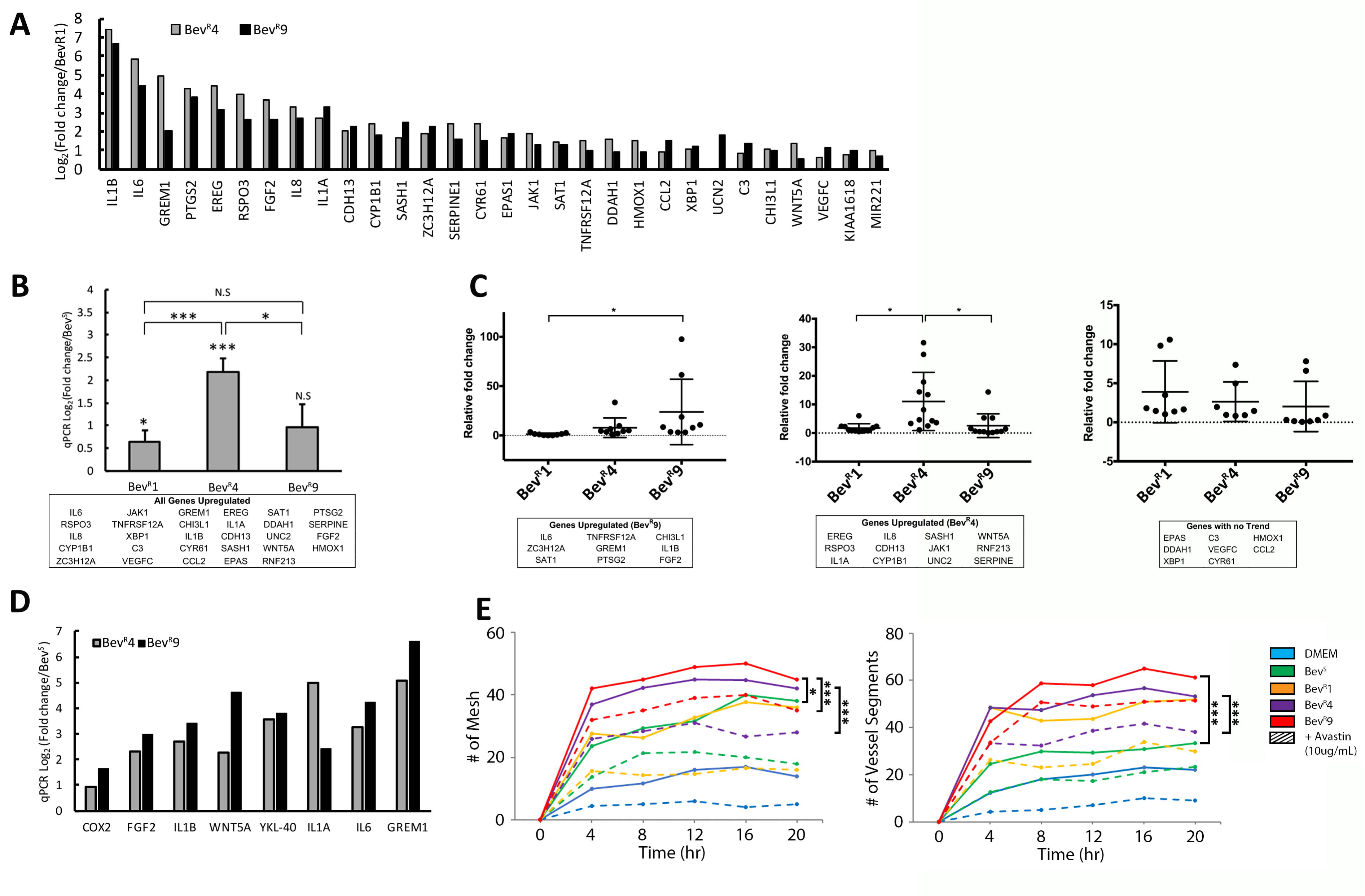
Increased but ineffective angiogenic gene expression in a multi-generational xenograft model of bevacizumab resistance. (**a**) The top 30 upregulated genes from the significantly enriched angiogenesis gene ontology were selected for validation by qPCR. (**b**) qPCR validation of 29 of the top 30 angiogenesis genes revealed significant overexpression of angiogenesis related genes by generation 1 of U87-Bev^R^, peaking by generation 4. (**c**) Dot plots separating the PCR results for the 29 pro-angiogenic genes into 3 categories: those that were upregulated with each U87-Bev^R^ generation, peaking at generation 9 (left); those that peaked in U87-Bev^R^ generation 4 (middle); and those with no trend from generation to generation of U87-Bev^R^ (right). (**d**) Multiple secreted VEGF-independent pro-angiogenic factors were upregulated across U87-Bev^R^ generations by qPCR. (**e**) HUVEC cells on matrigel were exposed to CM from U87-Bev^S^ and U87-Bev^R^ of multiple generations in the absence and presence of bevacizumab. U87-Bev^R^ CM drove increased endothelial segment and mesh formation compared to U87-Bev^S^ conditioned media (P=0.0007-0.01). While bevacizumab blocked U87-Bev^S^ effects on HUVEC cells (P=0.0004), by generation nine, bevacizumab failed to block the increased U87-Bev^R^ effects on HUVEC cells (P>0.05). *, p<0.05, **, p<0.01, ***, p<0.001

## DISCUSSION

The failure of anti-angiogenic agents like bevacizumab to achieve durable response in glioblastoma (5) and other cancers (21) has led to studies investigating the genetic reprogramming occurring in these tumors. Xenograft studies have suggested that tumor treatment with anti-VEGF antibody until tumor growth meets euthanasia criteria promotes increased transcription of mesenchymal factors that portend a worse prognosis in glioblastoma and other cancers (11, 12). However, this approach may not fully model patient resistance because it fails to replicate the duration of treatment patients receive, which can be far more prolonged than the duration of treatment that a xenograft receives before meeting euthanasia criteria. We sought to define the impact of prolonged duration of bevacizumab treatment on the genetic reprogramming of glioblastoma using our novel multi-generational xenograft model of bevacizumab resistance. The impact of this multi-generational approach was underscored by our finding that resistance did not become entrenched and irreversible until the fourth generation, the time at which the gene expression and functional changes associated with resistance peaked, suggesting that previous studies treating a single xenograft generation until euthanasia may not be using an adequate endpoint to model resistance to anti-angiogenic therapy.

Among the findings emerging from patient specimens and replicated from our xenograft model were the hypoxia and devascularization seen in BRGs, indicating that the resistance occurs despite continued successful devascularization caused by bevacizumab. Indeed, this devascularization succeeds despite upregulation of compensatory VEGF-independent pro-angiogenic pathways and despite enriched neurosphere yield in BRGs and bevacizumab-resistant xenografts and expression of genes associated with glioma stem cells, which have previously been shown to be capable of forming endothelial cells (22). This successful devascularization led to increased hypoxia in our resistance model, with an associated upregulation of genes involved in metabolism and inflammation, consistent with changes reported elsewhere (23-25).

Prognostic classification schemes have been created to designate molecular subtypes of glioblastomas based on similarity to defined expression signatures. Of the existing classification schemes, in order to study subtype changes associated with bevacizumab resistance, we chose to use that of Phillips et al. (14), which uses more focused gene sets to define subtypes that change during glioblastoma evolution, in contrast to other classification schemes like that of Verhaak et al. which uses more extensive gene sets to define glioblastoma subtypes that do not change during tumor evolution (26). Using this classification scheme, we demonstrated mesenchymal change in our multigenerational xenograft model which mirrored our finding of increased mesenchymal gene expression with longer duration of bevacizumab treatment in patient BRGs. In addition, we found that this mesenchymal change had an associated biomarker in the form of elevated serum YKL-40 in BRG patients. Similarly, serum YKL-40 elevation during bevacizumab treatment of ovarian cancer patients has been described as a predictor of shorter PFS (27) and a low baseline YKL-40 was associated with improved outcomes both in this ovarian cancer study (27) as well as among newly diagnosed glioblastoma patients receiving bevacizumab in a randomized trial (28).

Beyond the context of anti-angiogenic therapy, the present investigation adds to a growing number of studies implicating mesenchymal gene expression as a key hallmark of resistance to other anti-cancer therapies such as radiation therapy in glioblastoma (29) and EGFR inhibitors in lung cancer (30). These mesenchymal changes confer context-dependent advantages particular to the therapy that resistance is evolving against. In the case of anti-angiogenic therapy, we found that gene expression changes associated with a transition to a mesenchymal state conferred functional properties to tumor cells that would be advantageous in overcoming anti-angiogenic therapy. Using 3D bioengineered systems, we demonstrated a propensity for increased invasiveness, including perivascular invasiveness, in cells from our bevacizumab-resistant xenograft model. Invasiveness after bevacizumab resistance has often been described as perivascular, which supports tumor cells because the perivascular space is the entry point for nutrients into the brain (31-33). By invading alongside and engulfing preexisting cerebral microvasculature, perivascular invasion co-opts existing vasculature in a VEGF-independent and neo-angiogenesis-independent manner, providing a mechanism for continued GBM growth despite the tumor de-vascularization associated with VEGF blockade. Concomitant with this increased capacity for invasion, we also found that the gene expression changes during bevacizumab resistance promoted enrichment of the stem cell population, a progenitor cell type resistant to hypoxia and described as enriched in other studies of resistance to anti-angiogenic therapy (34).

Besides providing insight into potential biomarkers of bevacizumab resistance, our xenograft model could also offer insight into the optimal upstream regulators of bevacizumab resistance to target in order to prevent the evolution of resistance before it becomes entrenched. Because bevacizumab resistance has been challenging to pharmacologically target (35), these insights and the ability to screen inhibitors of bevacizumab resistance in our model could provide meaningful benefit to glioblastoma patients.

## MATERIALS AND METHODS

### Cell Culture

HUVEC cells (ATCC) were passaged fewer than 6 times, and confirmed to be Mycoplasma-free and verified by providing companies using short tandem repeat (STR) profiling. HUVEC cells were cultured in EGM-2 (Lonza). Cells for culture were obtained from U87-Bev^R^ and U87-Bev^S^ xenografts generated as described (10, 15). Cells were cultured in DMEM/F-12 supplemented with 10% FBS and 1% P/S and maintained at 37°C. U87-Bev^S^, U87-Bev^R^ and primary GBM neurospheres were cultured in Neurocult/Neurosphere media (StemCell Technologies) with 10 ng/ml bFGF (Thermo Fisher), 20 ng/ml EGF (Thermo Fisher) and B27 and N2 supplement (Thermo Fisher) and maintained at 37°C. Accutase (StemCell Technologies) was used to dissociate neurospheres into a single cell suspension.

### Animal Work

Animal experiments were approved by the UCSF IACUC (approval #AN105170-02). 500,000 U87 cells were implanted subcutaneously into SCID nude mice (3-4 weeks, female). Mice were then treated intraperitoneally with 10 mg/kg IgG control antibody (cat# I4506, Sigma) or bevacizumab (Genentech) twice weekly (n=15/group for generation one), with treating personnel blinded to treatment group. Tumor volumes were recorded twice weekly and bevacizumab resistance versus cure was defined by reaching endpoint specified by IACUC or complete tumor regression at 120 days. The largest IgG-or bevacizumab-treated tumors were harvested and dissociated into single cell suspensions which were then reimplanted subcutaneously into mice (500,000 cells per mouse; 10 mice/group for generations 2-9) after which treatment with bevacizumab or IgG commenced immediately and continued until size or symptomatic endpoint specified by IACUC criteria or tumor regression at 120 days. Resistance of the resulting U87-Bev^R^ and U87-Bev^S^ xenografts was confirmed in the orthotopic intracranial microenvironment at generations 6 and 9 as described previously (10). Tumor tissue from each generation was (1) dissociated into single cell suspensions for morphology analysis in culture; (2) perfused with PBS followed by 1% PFA, after which brains were isolated and kept in 4% PFA overnight, sunk in 30% sucrose, embedded in OCT, and cut on a cryostat for immunostaining; or (3) homogenized, then lysed into RIPA buffer for western blotting.

### Morphology Analysis

U87-Bev^S^ and U87-Bev^R^ (generations 1,4 and 9) were plated in a chamber slide system (Thermo Fisher) under normal cell culture conditions for 24 hours. Once adhered to the chamber slides, cells were fixed in 4% paraformaldehyde for 30 minutes, then washed three times with PBS after which cells were incubated in Phalloidin stain (1:200 in TNB; Sigma) for one hour at room temperature and washed again three times with PBS. Lastly, chamber slides were mounted with DAPI-Fluoromount G (Southern Biotech, 0100-20, Birmingham, AL, USA) and coverslipped. Slides were imaged using a Zeiss AxioObserver Z1 system equipped with an AxioCam MRm CCD and AxioVision software (Release 4.8). Cell morphology was quantified by calculating Form Factor [4πx area/√(perimeter)] using the Shape Descriptor plugin in ImageJ (NIH).

### Generation of Neurospheres and Functional Assays

Neurospheres were generated from patient GBM tumor by dissociating tumor chunks into a single cell suspension in full Neurocult media at a density of 100,000 cells. U87-Bev^S^ and U87-Bev^R^ neurospheres were generated by seeding the respective cell lines at 50,000, 100,000 and 200,000 cells in neurosphere media. Neurospheres were isolated and stained for cancer stem cell markers (CD133 and Nestin) as described by Kim et al. (36) to confirm stem cell isolation. Neurosphere reformation assay, diameter quantification and total cell counts were done by seeding 100,000 cancer stem cells in neurosphere media. At the end of 7 days, tile imaging was performed using a Zeiss AxioObserver Z1 system from which number of spheres and diameters were calculated. Single cell suspensions were prepared from the neurospheres to get absolute cell counts from patient GBM neurospheres as well as U87-Bev^S^ and Bev^R^ cell lines.

### CRISPR knockout

ZEB1 was knocked out in U87-Bev^S^, U87-Bev^R^ generations 1,4 and 9 by co-transfecting ZEB1 CRISPR/Cas9 KO (sc-400201; Santa Cruz Biotechnology, CA) and ZEB1 HDR Plasmids (sc-400201-HDR; Santa Cruz Biotechnology, CA) using FuGENE® 6 Transfection Reagent (Promega). Transfections were done as per manufacturer instructions. Successful transfections were confirmed using qPCR and fluorescent microscopy for Red Fluorescent Protein (RFP).

### Microarray Analysis

Tumor chunks previously harvested from generational xenografts had been flash frozen at explantation and stored in liquid nitrogen. Samples for the microarray analysis were retrieved from storage and dissociated using both a QiaShredder (Qiagen) and passage through a 21-gauge sterile syringe. Dissociated tissue was then processed to obtain RNA using the RNeasy kit (Qiagen), following standard manufacturer’s protocol. RNA was tested for quality using an RNA 6000 chip with the Bioanlyzer hardware (Agilent). RIN scores of greater than 8 were required to ensure RNA quality. RNA was then converted to labeled cRNA using the TargetAmp-Nano Labelling Kit for Illumina Expression BeadChip (EpiCentre), following standard manufacturer’s protocol. Labelled cRNA was kept at −20°C and given to the UCSF Genome Core Facility (GCF) for chip hybridization.

### Single-cell RNA sequencing

Tissue was dissociated by incubation in papain with 10% DNAse for 30 min. A single-cell suspension was obtained by manual trituration using a glass pipette. The cells were filtered via an ovomucoid gradient to remove debris, pelleted, and resuspended in Neural Basal Media with serum at a concentration of 1700 cells/μL. In total, 10.2 μL of cells were loaded into each well of a 10X Chromium Single Cell capture chip and a total of two lanes were captured. Single-cell capture, reverse transcription, cell lysis, and library preparation were performed per manufacturer’s protocol. Sequencing was performed on a HiSeq 2500 (Illumina, 100-bp paired-end protocol). Raw data was processed with CellRangeR 1.3. The resulting count table derived from CellRangeR was processed in R 3.4.1. Data normalization (log (CPM/100+1) and subsequent tsne clustering was performed with the Seurat R package (37). Copy number inference was carried out with the CONICSmat R package (https://github.com/diazlab/CONICS/). Only tumor cells harboring at least one clonal mutation (chr10 loss or chr7 gain, determined by thresholding the posterior probability of the mixture model, pp<0.05) were taken into account for tsne clustering.

### Bioinformatics

Data (.idat files) from GCF analysis of our xenograft arrays underwent standard QC and processing through the UCSF Bioinformatics Core, and were deposited in GEO (Accession number=GSE81465). In parallel, we accessed our previously archived (13) microarray analysis of BRGs and their paired pre-treatment GBMs deposited in ArrayExpress (accession no. E-MEXP-3296). Gene set enrichment analysis was performed in R using the GAGE package (v2.28.2) (38). A pre-existing set 1000 gene ontology (GO) gene lists within the GAGE package was used to identify enriched gene ontologies in the positive (upregulated) and negative (downregulated) directions. To identify significantly dysregulated genes across generations, a two-component normal mixture-model was fitted by expectation-maximization to the Z-transformed Log(variance) distribution of gene expression values over generations 4 and 9. A posterior probability cutoff of ≥0.95 was applied, resulting in 717 significantly dysregulated genes (39). Next, a non-parametric bootstrap procedure was used to determine over-representation of the Philips et al mesenchymal, proneural, and proliferative gene sets among these dysregulated genes. Microarray probes were mapped to HGNC gene names, and uniform random sampling of 717 of the 48,324 gene probes followed by matching to the mesenchymal, proneural, or proliferative gene sets was repeated 50,000 times, resulting in a putative null distribution of the number of matches to the gene set in question. A Poisson probability density function was fit by maximum likelihood estimation to this distribution and a 1-tailed p-value calculated for the observed number of matches to the gene set in question. Genes were further clustered by k-means clustering based on gene expression trajectory across generations, and these expression trajectory clusters were manually curated into sets of upregulated genes (sustained increases in expression through generation 4 and 9), downregulated genes (sustained decreases in expression through generation 4 and 9), and other genes. Finally, expression of particular genes of interest identified from microarray analysis were validated using qPCR.

### Quantitative PCR

After obtaining RNA from xenografts as described in the microarray analysis, cDNA was created using qScript XLT cDNA Supermix (Quanta Bio), both following standard manufacturer’s protocol. cDNA was diluted to a constant concentration for all samples to ensure similar nucleic acid loading levels. Quantitative PCR was carried out using Power Syber Green Master Mix (Applied Biosystems). Primer sequences can be found in **Supplementary Tables S1-S2**. qPCR was performed on an Applied Biosystems StepOne Real-Time PCR cycler following recommended guidelines described by Applied Biosystems for Syber: 95**°** C for 10 min, followed by 40 cycles of 95**°** C for 15 sec and 60**°** C for 1 min. Ct values were calculated using the StepOne software accompanying the real-time cycler. Samples were prepared with three technical replicates for each primer pair and used ACTB as a control housekeeping gene.

### Human Serum ELISA

GBM patient serum was obtained through the UCSF Brain Tumor tissue bank which, through an IRB-approved protocol (10-01318), obtained tissue after informed consent from patients. YKL-40 ELISAs (R&D Systems) were performed in triplicate on serums diluted 1:50.

### Microchannel Device Fabrication

The microchannel device was fabricated as described by Lin et al. (16). Briefly, the micropatterned polyacrylamide (PA) gel (8%T, 3.3%C acrylamide/bisacrylamide (Sigma-Aldrich) was activated with Sulfo-SANPAH (Thermo Scientific) and incubated with a 1:2 dilution of Matrigel (Fisher Scientific, Cat# CB40230, diluted in 10% FBS cell culture medium) for 24 hours at 4°C (see **Supplementary Methods**). The micropatterned PA gel was then incubated in 0% FBS cell culture medium at 37°C for at least 1 hour prior to device fabrication. Following incubation, the gel was removed and the excess liquid was aspirated away. The PDMS lid was gently placed on top of the PA gel (see **Supplementary Methods**), with the holes in the PDMS aligned to the inlet and outlet microwell patterns on the PA gel. The hybrid PDMS/PA device was then placed under vacuum (200 mmHg) in a vacuum desiccator (Bel-Art) for 6 minutes to reversibly seal the layers together. After the vacuum step, 0% FBS cell culture media was place in both inlet ports and allowed to equilibrate in the device. To seed the device, any excess medium was first aspirated from all four inlet and outlet ports. Then a 30 µl aliquot of cell suspension (1 million cells per mL) was added to the top inlet port. Due to the pressure differences, cells flow into and lodge at the start of the migratory microchannels. The cells were then allowed to adhere for one hour prior to chemokine gradient formation and imaging, with 20 ng/ml SDF-1 (R&D Systems) diluted in 10% FBS used as the chemokine.

### Cell Motility Measurements in Microchannel Device

Live-cell imaging was performed using a Nikon Ti-E2000-E2 microscope equipped with a motorized, programmable stage (Applied Scientific Instrumentation), an incubator chamber to maintain constant temperature, humidity, and CO2 levels (In vivo Scientific), a digital camera (Photometrics CoolSNAP HG II), and NIS Elements (Nikon Instruments Inc.) software. Images were taken at 5 ms exposure, 2×2 pixel binning using a 10x-objective (Nikon CFI Plan Fluor DLL 10x). We measured cellular motility using 10X phase contrast time-lapse images acquired every 15 minutes over a 3-hour period. ImageJ software (NIH) was used to track the centroid of each cell from one frame to another to yield instantaneous migration speeds, which were then averaged over the entire time course of the experiment to yield the migration speed of a cell. Cells that were observed to be sticking to each other were also excluded from analysis.

### HA Matrix Synthesis

HA hydrogels were synthesized as previously described (40). Briefly, sodium hyaluronate (Lifecore Biomedical, Research Grade, 66 kDa −99 kDa) was functionalized with methacrylate groups (Me-HA) using methacrylic anhydride (Sigma-Aldrich, 94%). The extent of methacrylation per disaccharide was quantified by ^1^H NMR as detailed previously and found to be ~85% for materials used in this study. To add integrin-adhesive functionality, Me-HA was conjugated via Michael Addition with the cysteine-containing RGD peptide Ac-GCGYGRGDSPG-NH2 (Anaspec) at a concentration of 0.5 mmol/L. Finally, 3wt% Me-HA was crosslinked in phenol-free DMEM (Invitrogen) with bifunctional thiol dithiothretiol (DTT, Sigma-Aldrich). After 1 hr of crosslinking, the hydrogels were rinsed and soaked in room temperature PBS for 1 hr before cell seeding. The shear modulus of hydrogel formulations was measured using oscillatory rheometry (Anton Parr Physica MCR 310) as described previously.^25^ Briefly, hydrogels were first crosslinked by incubation for 1 hr in a humidified 37˚C chamber. Rheological testing consisted of frequency sweeps ranging from 100 to 0.1 Hz at 0.5% amplitude also in a humidified 37˚C chamber. Shear modulus was reported as the average storage modulus for 3 tests per type of matrix composition at an oscillation frequency of 0.5 Hz. A concentration of 19 mmol/L DTT was selected to yield a shear modulus of ~300 kPa for all studies.

### HA Invasion Device Fabrication

Device fabrication will be described in detail in a forthcoming manuscript. Briefly, the PDMS PDMS (Sylgard 184 elastomer) supports for hydrogels were fabricated by molding PDMS to have two open channels (~170 µm in diameter) in parallel with ~500 µm spacing between channels. A gap was cut into the PDMS allowing space for gel casting. To cast the gel, wires were first reinserted into the assembled device to create a mold. Next, the hydrogel matrix was inserted in the side of the device. Wires were removed after the hydrogel solution had solidified leaving two open channels. After rinsing and soaking the hydrogels, cells were inserted into the freshly fabricated device using a syringe (Hamilton). Cut wires were used to plug each end of the cell reservoir, and the entire device was placed into the bottom of a 6-well plate and bathed in 3 mL of medium. Cells and gels were equilibrated in medium overnight before imaging. Medium was changed every 3 days.

### Invasion and Protrusion Analysis

For area analysis in HA Invasion devices, cells in devices were imaged once every 3 days using Eclipse TE2000 Nikon Microscope with a Plan Fluor Ph1 10x objective. Images were acquired and large images were stitched using NIS-Elements Software. For each device, the total cell area was outlined in ImageJ and normalized to the total cell area from day 1. To analyze morphology, protrusions visible at the edge of the invading mass of cells were counted and normalized to the length of the cell mass. Each device was fabricated and seeded independently. GraphPad Prism 7 software was used to graph data and perform statistical analysis. Data was compared using ANOVA followed by Tukey Kramer multiple comparisons test (n=3-4 independent devices per condition).

### Blood Vessel Formation Assay

U87-Bev^R^ or U87-Bev^S^ cells from variable generations were grown in DMEM/10% FBS in the upper reservoir of Boyden chambers coated with 30 μg/mL Collagen I (Life Technologies) in PBS for 24 hours. On day 2, HUVEC cells were seeded over matrigel (Corning) in the lower reservoir of each Boyden chamber in media with or without 10μg/mL bevacizumab. Thirty minutes prior to imaging, calcein-AM (Life Technologies) was added at a final concentration of 2 μg/mL to each lower reservoir. Cells were imaged every 4 hours for 20 hours. The ImageJ plug-in Angiogenesis Analyzer was used to compute number of mesh and vessel segments, as described previously (41).

### Statistics

The student’s t-test and Mann-Whitney tests were used to compare means of continuous parametric and non-parametric variables, respectively. Paired t-tests (parametric) and Wilcoxon signed-rank tests (non-parametric) were used to compare pre-and post-treatment variables from the same patients. Interobserver variability for manual counting of immunofluorescence was assessed using SPSS VARCOMP analysis. *P* values are two-tailed and *P*<0.05 was considered significant.

## CONFLICT OF INTEREST

The authors report no competing financial interests in relation to the work described.

## ACKNOWLEDGEMENTS

M.K.A. was supported by the American Brain Tumor Association (ABTA), the James S. McDonnell Foundation, American Cancer Society (ACS), University of California Cancer Research Coordinating Committee, and the NIH (5K02NS64167 and 1R01NS079697). A.J., A.C., and J.R. were supported by Howard Hughes Medical Institute (HHMI) fellowships. A.J. was supported by an NIH Predoctoral Fellowship (1F31CA203372-01). W.C. and J.R. were supported by UCSF School of Medicine (SOM) Pathways Explore Summer Grants. K.J.W. was sponsored by an NSF Predoctoral Fellowship. S.K. was sponsored by the NIH (1R21CA174573, 1R01GM122375, 1R21EB025017).

## Notes

The authors have declared that no conflict of interest exists.

